# CDR Proteins of *C*. *elegans* and Related Species: Relationship to Metaxin and FAXC Proteins

**DOI:** 10.1101/2024.01.02.573863

**Authors:** Kenneth W. Adolph

## Abstract

CDR (cadmium-responsive) proteins of the nematode *C. elegans* and related species are shown to have structural homology to metaxin proteins and FAXC (failed axon connections) proteins of vertebrates and invertebrates. Unlike the metaxin and FAXC proteins, however, the predicted CDR proteins are only encoded in the genomes of nematodes, and not in vertebrates or other invertebrates. Metaxin-like structural features of CDR proteins are shown in this report to include: (1) GST_N_Metaxin and GST_C_Metaxin conserved protein domains, (2) metaxin-like patterns of α-helical secondary structure, and (3) a special 4-stranded β-sheet motif shared with the metaxins. FAXC proteins also possess these structural features. Phylogenetic analysis revealed that CDR proteins are related to metaxins 1, 2, and 3 and to FAXC proteins, although more closely to FAXC proteins. Nevertheless, all three types of proteins – CDR, FAXC, and metaxin – are related by evolution. Although CDR proteins have structural homology to metaxin and FAXC proteins, pairwise alignments of CDR proteins with metaxins and FAXCs demonstrated only low percentages of identical amino acids. CDR proteins can therefore be considered a separate category of proteins, distinct from metaxin and FAXC proteins. The CDR genes of *C. elegans* consist of a family of seven genes, *cdr-1* to *cdr-7*, which form a cluster of closely adjacent genes on chromosome V. Multiple genes were also found for the related species *C. briggsae* and *C. remanei*. Alignment of pairs of *C. elegans* CDR protein sequences encoded by different *cdr* genes demonstrated a high level of sequence homology. The neighboring genes of the *C*. *elegans cdr* genes are different than the genes adjacent to the *C*. *elegans faxc* genes and the *mtx-1* and *mtx-2* metaxin genes.

## 1. INTRODUCTION

The cadmium-responsive gene *cdr-1* was initially described as representing a new class of cadmium-inducible genes in *C. elegans* nematodes (Liao et al., 2002). The CDR-1 protein was classified as an integral membrane protein of lysosomes that protects against the toxicity of the heavy metal cadmium. A family of *C. elegans* genes was identified with genes that are homologous to *cdr-1* (Dong et al., 2005). These include *cdr-2* to *cdr-7*, and, like *cdr-1*, appear to encode integral membrane proteins. Deletion of the *cdr-1* gene did not lead to hypersensitivity to cadmium toxicity, due to an increase in heavy metal chelators (Hall et al., 2012).

As demonstrated in this paper, CDR proteins possess the same structural features that characterize the metaxins, even though CDR proteins and metaxins are different proteins. The metaxins, originally described in vertebrates, are mitochondrial membrane proteins that function in the uptake of proteins into mitochondria. The structural features of CDR proteins were examined in comparison with metaxin proteins and the metaxin-like FAXC proteins. CDR proteins were found to have metaxin-like features that include the major conserved protein domains of the metaxins (GST_N_Metaxin and GST_C_Metaxin domains). In addition, CDR proteins, like metaxin proteins, have secondary structures dominated by patterns of α-helical segments. And the 3D structures of CDR proteins have a special 4-stranded β-sheet motif that is characteristic of the metaxins. FAXC proteins, first described in *Drosophila melanogaster* (Hill et al., 1995), also have the same structural features, with conserved GST_N_ and GST_C_Metaxin domains, α-helical secondary structures, and the characteristic β-sheet motif. Both metaxin and FAXC genes are found in a wide variety of organisms, unlike *cdr* genes, which are confined to nematodes.

Three metaxins (1, 2, and 3) have been revealed in vertebrates including humans and also the mouse (genome sequence: Mouse Genome Sequencing Consortium, 2002), frog *Xenopus laevis* (Session et al., 2016), and zebrafish *Danio rerio* (Howe et al., 2013). Moreover, metaxin-like proteins have been detected in invertebrates (Adolph, 2020a), plants and bacteria (Adolph, 2020b), and protists and fungi (Adolph, 2021).

FAXC (failed axon connections) proteins are also members of the family of metaxin-like proteins (Adolph, 2023). They are encoded in the genomes of vertebrates and a variety of invertebrates. Invertebrate phyla that possess genes for FAXC proteins include Arthropoda, Mollusca, Cnidaria, and Placozoa. Structural features of the predicted FAXC proteins include the same protein domains and α-helical and β-sheet secondary structures that define the metaxins and metaxin-like proteins.

The *C. elegans* roundworm has been extensively used as a model research organism, especially in studies of neural development and other areas of molecular and developmental biology. For example, the pattern of neurons has been mapped in detail to produce a “connectome” of neuronal interactions. The developmental fate of every somatic cell has been mapped to determine the patterns of cell lineage. The roundworm has also been useful in studies of programmed cell death, ageing, meiosis, and a variety of other biomedical research topics. Further, the *C. elegans* genome was the first genome of a multicellular organism to be fully sequenced (*C. elegans* Sequencing Consortium, 1998).

Very little has been reported in the literature regarding *cdr* genes and CDR proteins beyond the first description of the *cdr-1* gene and follow-up studies concerned with the family of *cdr* genes, the role of *cdr-1* in cadmium detoxification, and related aspects. For the study reported here, initial searches of NCBI databases revealed that CDR proteins have homology to metaxin and FAXC proteins. This investigation was therefore undertaken to determine more fully the properties of CDR proteins and the relationship between CDR, metaxin, and FAXC proteins. The results support the conclusion that CDR proteins are members of the metaxin-like family of proteins. In particular, the proteins have metaxin-like properties including GST_N_Metaxin and GST_C_Metaxin domains, metaxin-like patterns of α-helical secondary structure, and a characteristic 3D β-sheet motif also found with metaxin proteins. Much more needs to be understood, however, about the functions of these proteins, which further studies might reveal.

## 2. METHODS

Protein domain structures were investigated using the NCBI CD search tool (www.ncbi.nlm.nih.gov/Structure/cdd/wrpsb.cgi; Lu et al., 2020). Figure 1 shows the major domains of CDR, metaxin, and FAXC proteins. α-Helical and β-strand secondary structures were predicted with the PSIPRED server (<u>bioinf.cs.ucl.ac.uk/psipred/</u>; Jones, 1999; Buchan and Jones, 2019). The COBALT multiple-sequence alignment tool (www.ncbi.nlm.nih.gov/tools/cobalt; Papadopoulos and Agarwala, 2007) was used to produce the alignments of α-helical and β-strand segments in Figure 2. Transmembrane α-helices were identified with PHOBIUS (Kall et al., 2007) and TMHMM (Krogh et al., 2001). The AlphaFold protein structure database (Jumper et al., 2021; Varadi et al., 2022) was employed to investigate the predicted 3D structures of the proteins, including the characteristic 3D β-sheet motif (Figure 3). The AlphaFold structures were viewed through NCBI BLAST searches and the iCn3d structure viewer (Wang et al., 2020). The phylogenetic relationships of CDR, metaxin, and FAXC proteins in Figure 4 were also derived from multiple-sequence alignments produced by the COBALT alignment tool. In addition, Clustal OMEGA (https://www.ebi.ac.uk/Tools/msa/clustalo/) was used to generate phylogenetic trees. Identical and similar amino acids of pairs of protein sequences (Figure 5 and Figure 7) were determined with the NCBI BLAST Global Align tool and the Align Two Sequences tool (https://blast.ncbi.nlm.nih.gov/Blast.cgi; Needleman and Wunsch, 1970; Altschul et al., 1990).

**Figure 1.**
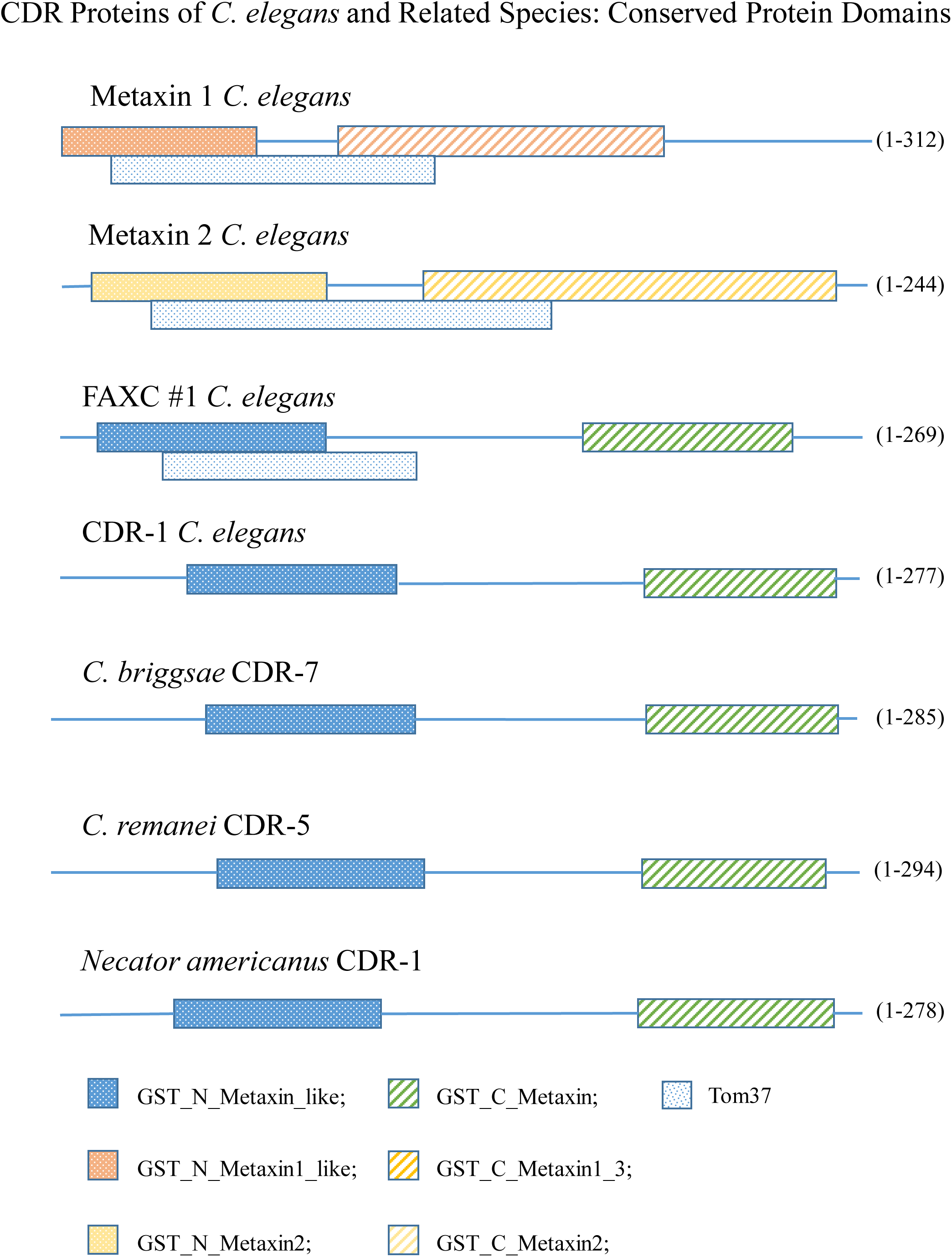
Conserved GST_N_Metaxin and GST_C_Metaxin domains of CDR proteins. The figure includes the major conserved domains of CDR proteins of *C. elegans* and the closely related nematodes *C. briggsae* and *C. remanei*, and also *Necator americanus*. These domains are characteristic of the metaxins and FAXC proteins in addition to CDR proteins. This is seen in the domains of *C. elegans* metaxin 1, metaxin 2, and FAXC #1. The Tom37 domain is also a common protein domain of metaxin and FAXC proteins, as shown for metaxins 1 and 2 and FAXC #1 of *C. elegans*. However, Tom37 is not a distinguishing domain of CDR proteins. All of the multiple CDR proteins of *C. elegans* (CDR-1 to CDR-7) have the same conserved GST_N_Metaxin and GST_C_Metaxin domains. *C. briggsa*e and *C. remanei* also have multiple CDR proteins with these domains.

**Figure 2.**
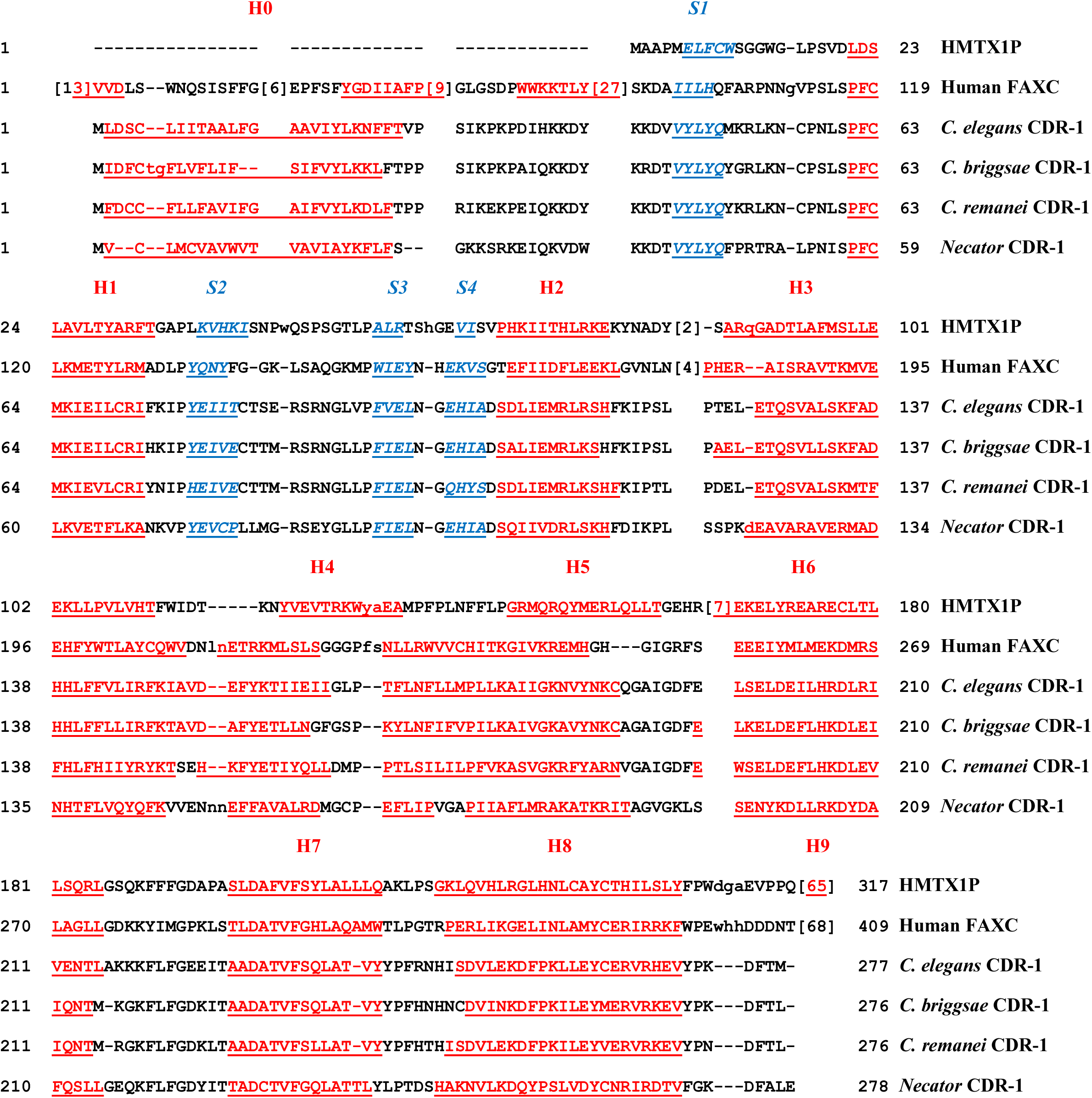
Alpha-helical and β-strand secondary structure of CDR proteins. The figure demonstrates that α-helical segments are the dominant feature of the secondary structure of CDR proteins, but with significant, conserved β-strand segments. In the figure, the α-helical sequences are in red and are underlined. β-strand segments are in blue, and are italicized and underlined. The primary helical segments are labeled H1 through H8 based on the labeling for the metaxins. The β-strand segments are labeled S1 through S4. The CDR-1 proteins have an extra helix H0 at the N-terminus, while metaxin 1 has an extra helix H9 at the C-terminus (within [65] in the figure). The multiple sequence alignment includes the CDR-1 proteins of *C. elegans*, *C. briggsae*, *C. remanei*, and *Necator americanus*. The patterns of α-helical segments and β-strand segments are highly conserved among the CDR-1 proteins. Furthermore, the patterns of secondary structure for the CDR proteins are nearly identical to the patterns for human metaxin 1 (HMTX1P) and human FAXC isoform 1, except for differences near the N- and C- termini.

**Figure 3.**
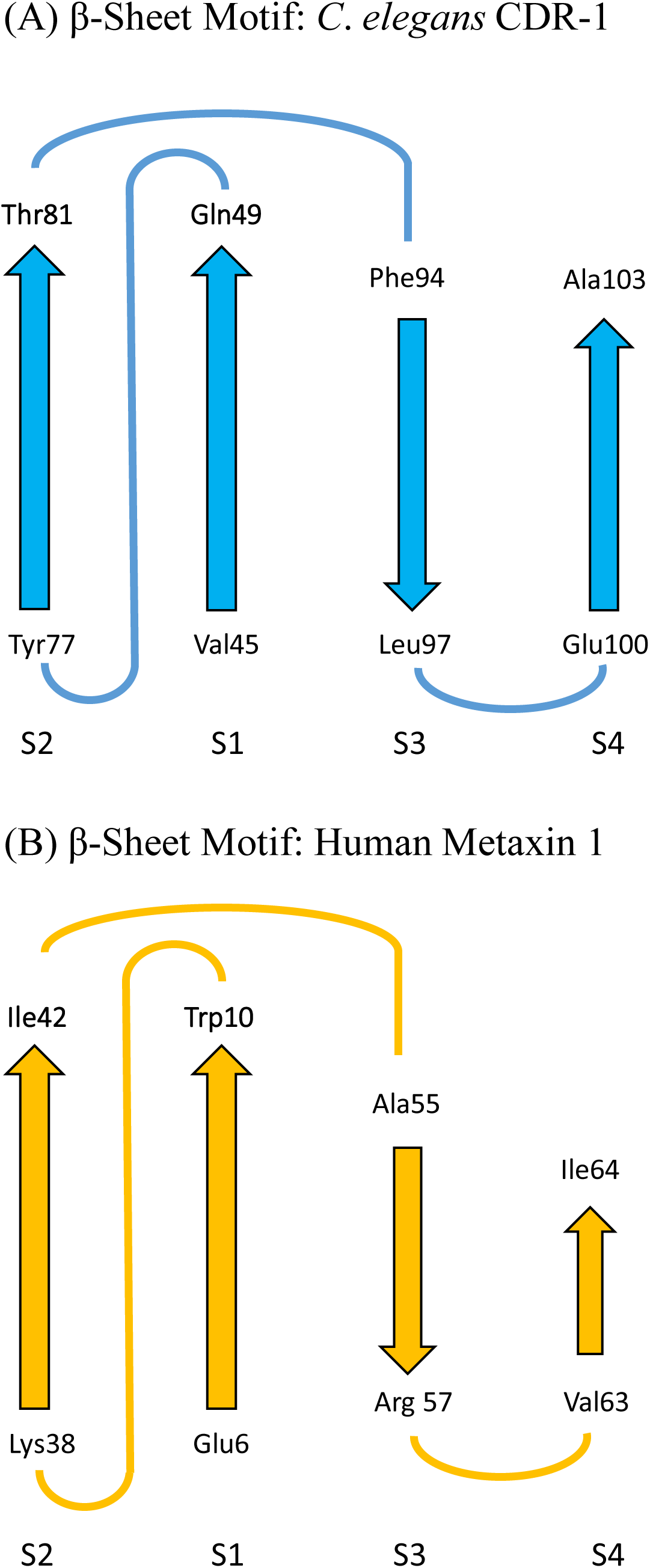
Conserved 3D β-sheet motif of CDR, metaxin, and FAXC proteins. A characteristic 4-stranded β-sheet structure, shown in Figure 3A and 3B, was revealed in the predicted 3D structures provided by AlphaFold (Jumper et al., 2021). The β-sheet motif was detected in CDR proteins and also in the more widely distributed metaxin and FAXC proteins. Figure 3A includes the motif for *C*. *elegans* CDR-1. The same motif was present in other CDRs of *C*. *elegans* and different nematodes. The strands are in the order S2, S1, S3, S4, and the orientation of the strands is **↑ ↑ ↓ ↑**. The arrangement of the strands for human metaxin 1 is shown in Figure 3B. For both (A) and (B), the N-terminal and C-terminal amino acids are indicated for each strand. Although the β-strands in (A) and (B) are relatively short and show some variation in length, the order and orientation of the strands are the same. The conserved β- sheet motif is a characteristic feature of the 3D structures of other vertebrate and invertebrate metaxin proteins and FAXC proteins, including vertebrate metaxins 1, 2, and 3 and invertebrate metaxins 1 and 2. The metaxin and FAXC proteins of fungi and the FAXC proteins of bacteria also possess the motif.

**Figure 4.**
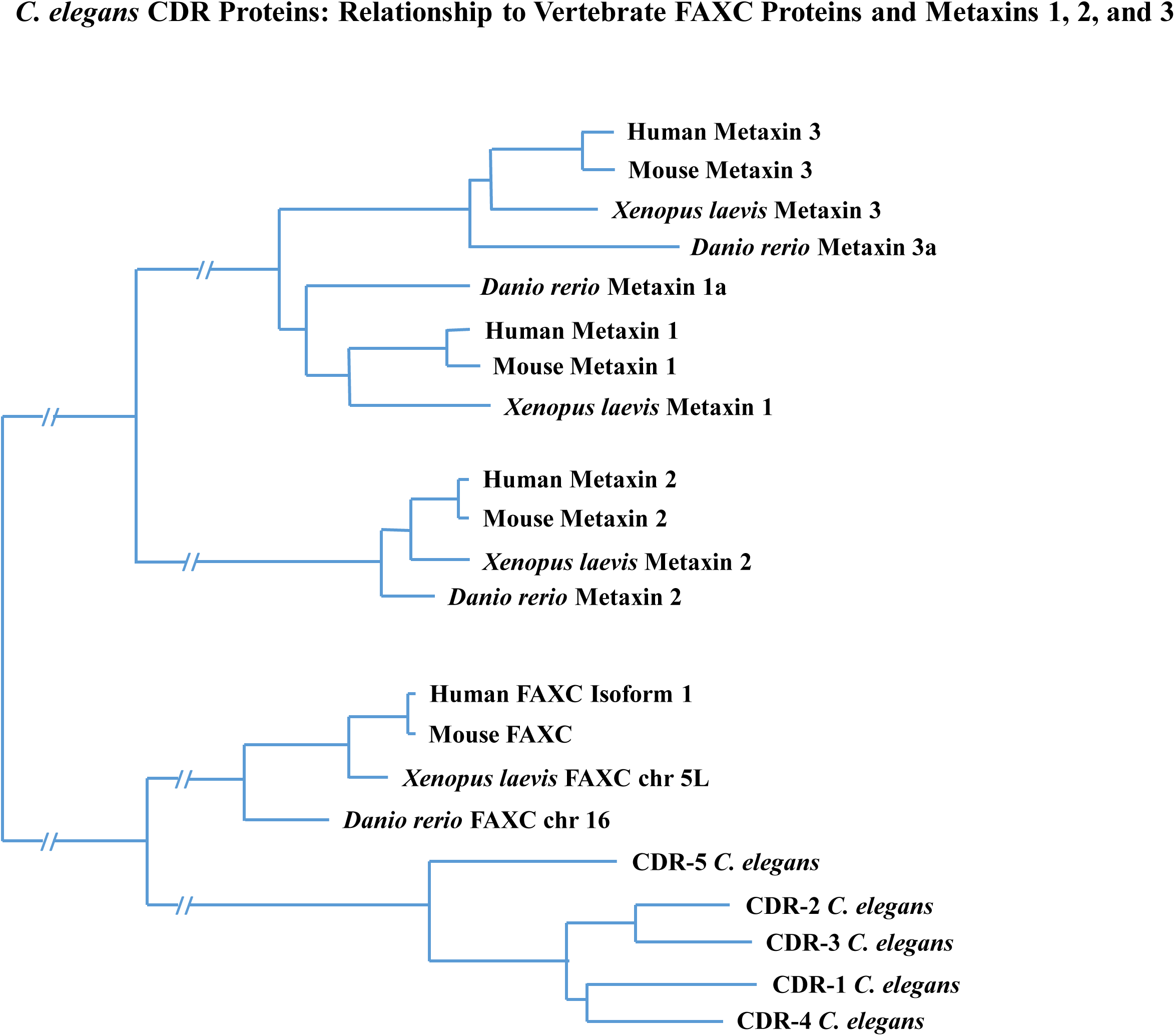
Phylogenetic relationships of CDR, FAXC, and metaxin proteins. The analysis of evolutionary relationships as shown in the figure reveals that CDR proteins form a separate group compared to the FAXC and metaxin proteins. Most significantly, the CDR and FAXC proteins, and the metaxin 1, metaxin 2, and metaxin 3 proteins, are all related by evolution. These phylogenetic results are in accord with the conserved structural features (protein domains, α-helices, β-strand segments) of the proteins. Figure 4 includes CDR-1 through CDR-5 of *C. elegans*, and shows that CDR proteins are more similar to the FAXC group of proteins than to the metaxin 1, 2, and 3 groups. The most closely related metaxins are metaxins 1 and 3, while the metaxin 2 proteins are less related. In the figure, the lengths of the branches are proportional to the degree of evolutionary change of the different proteins.

**Figure 5.**
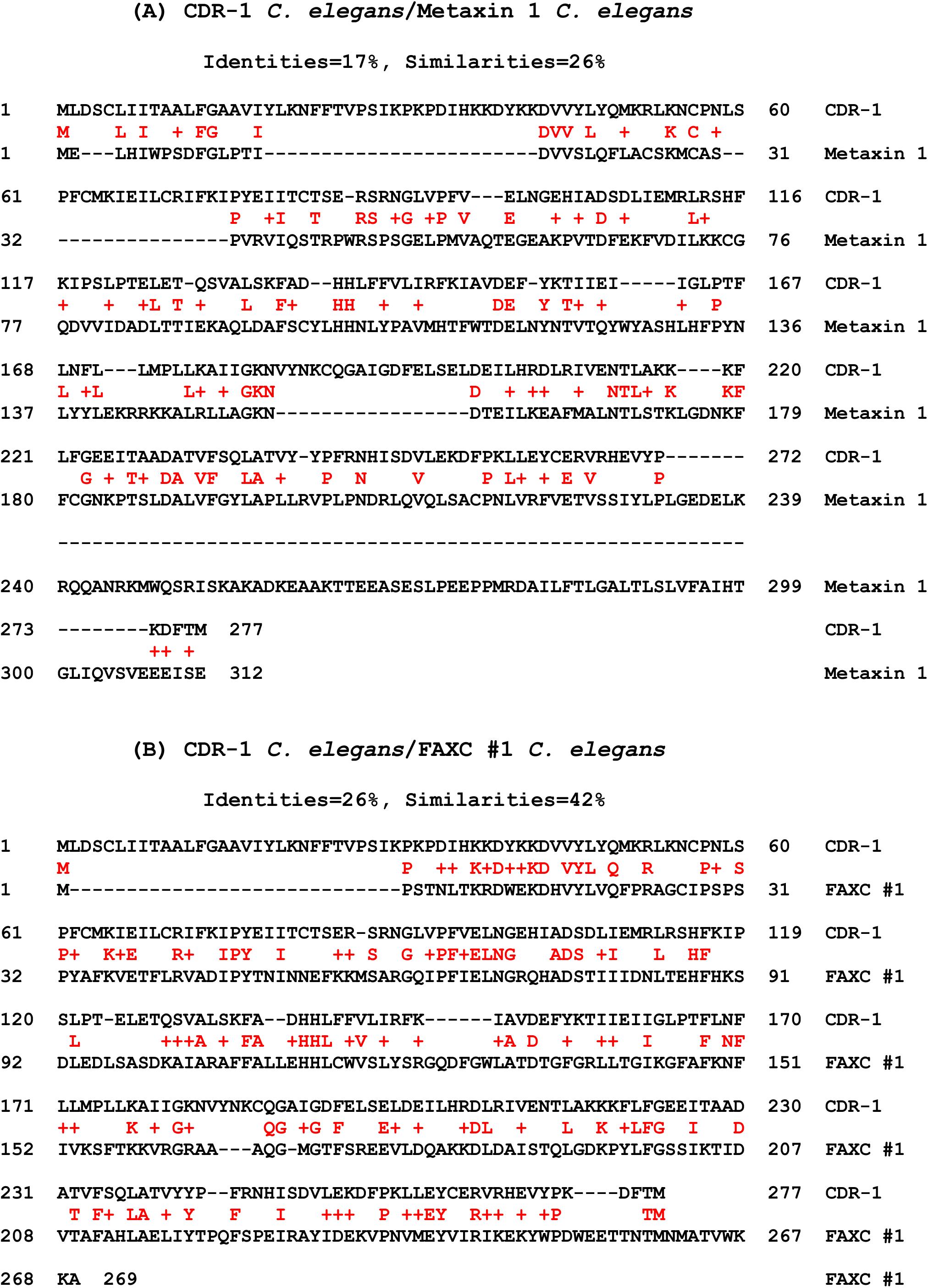
Alignment of CDR protein sequences with metaxin and FAXC sequences. Only low percentages of identical amino acids are found in comparing CDR proteins with metaxin and FAXC proteins. In Figure 5A, CDR-1 is aligned with metaxin 1 (both *C. elegans*). Low percentages of identical amino acids (17%) and similar amino acids (26%) are detected. In (B), CDR-1 and FAXC #1 of *C. elegans* are aligned, and also display relatively low percentages of identical (26%) and similar (42%) amino acids. The alignments in (A) and (B) used NCBI Global Align, which compares amino acid sequences along their whole lengths. Other combinations of CDR proteins (CDR-2 through CDR-7) with different metaxin and FAXC proteins provided comparable results. The general conclusion from these alignments is that CDR, metaxin, and FAXC proteins constitute different protein categories. This is the case even though the proteins have similar structural features, in particular similar conserved protein domains and secondary structures.

The EMBOSS tool (https://www.ebi.ac.uk/Tools/psa/emboss_needle/) was also utilized to align amino acid sequences. Genomic regions of *Caenorhabditis* species with multiple, adjacent *cdr* genes (Figure 6) were identified using the NCBI “Gene” database (www.ncbi.nlm.nih.gov/gene).

**Figure 6.**
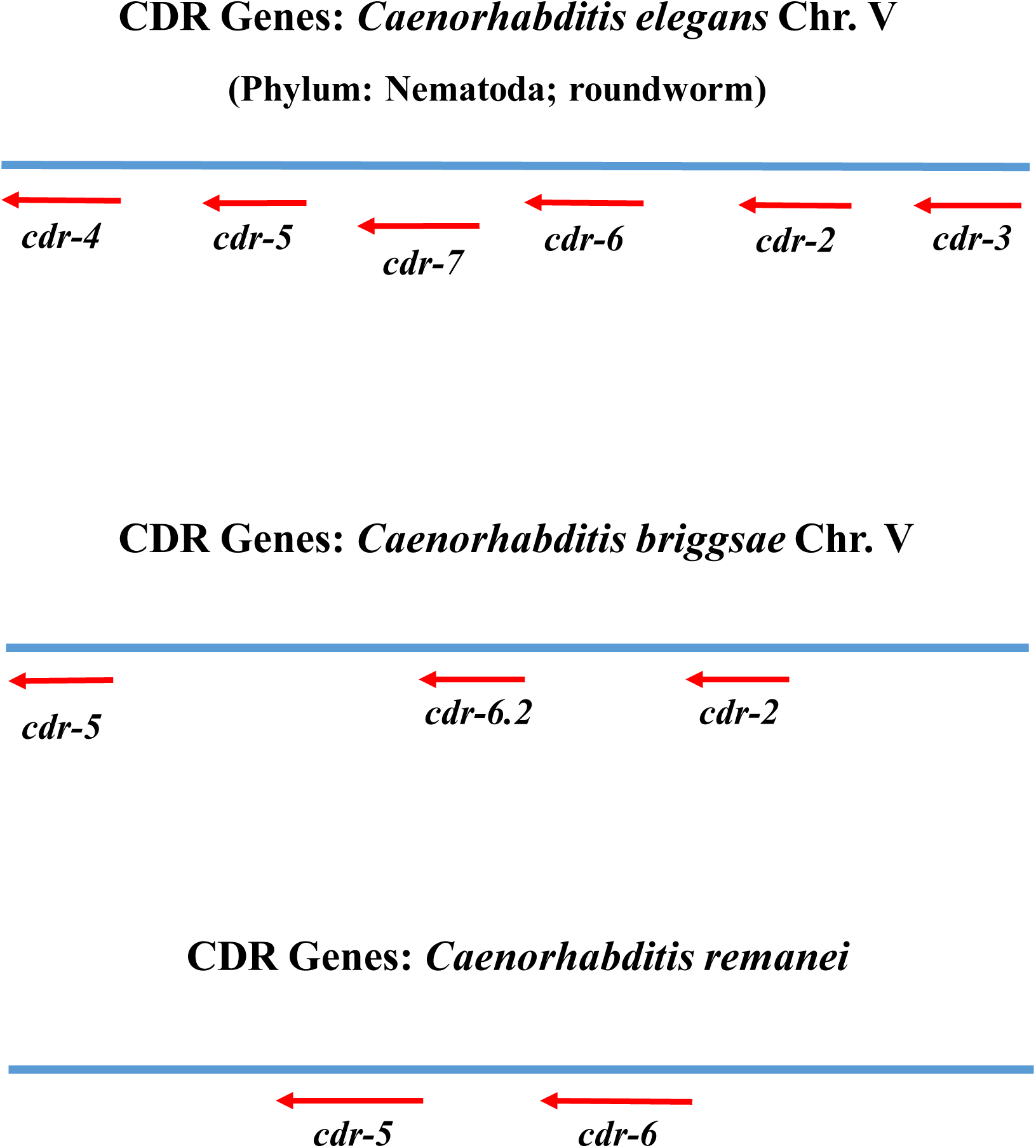
Genomic regions of the multiple CDR genes of *C. elegans* and related nematode species. The figure includes the genomic regions of *C. elegans* (top of the figure), *C. briggsae* (middle), and *C. remanei* (bottom) that have multiple CDR genes in close proximity. For *C. elegans*, six of the seven identified CDR genes (*cdr-2* to cdr-7) are clustered in the genomic region, which is on chromosome V. The genome of *C. elegans* is made up of five autosomes (I – V) and an X chromosome. The *C. briggsae* region, on *C. briggsae* chromosome V, includes three CDR genes (*cdr-2*, *cdr-5*, and *cdr-6.2*). For *C. remanei*, two genes (*cdr-5* and *cdr-6*) are found in the genomic region shown.

**Figure 7.**
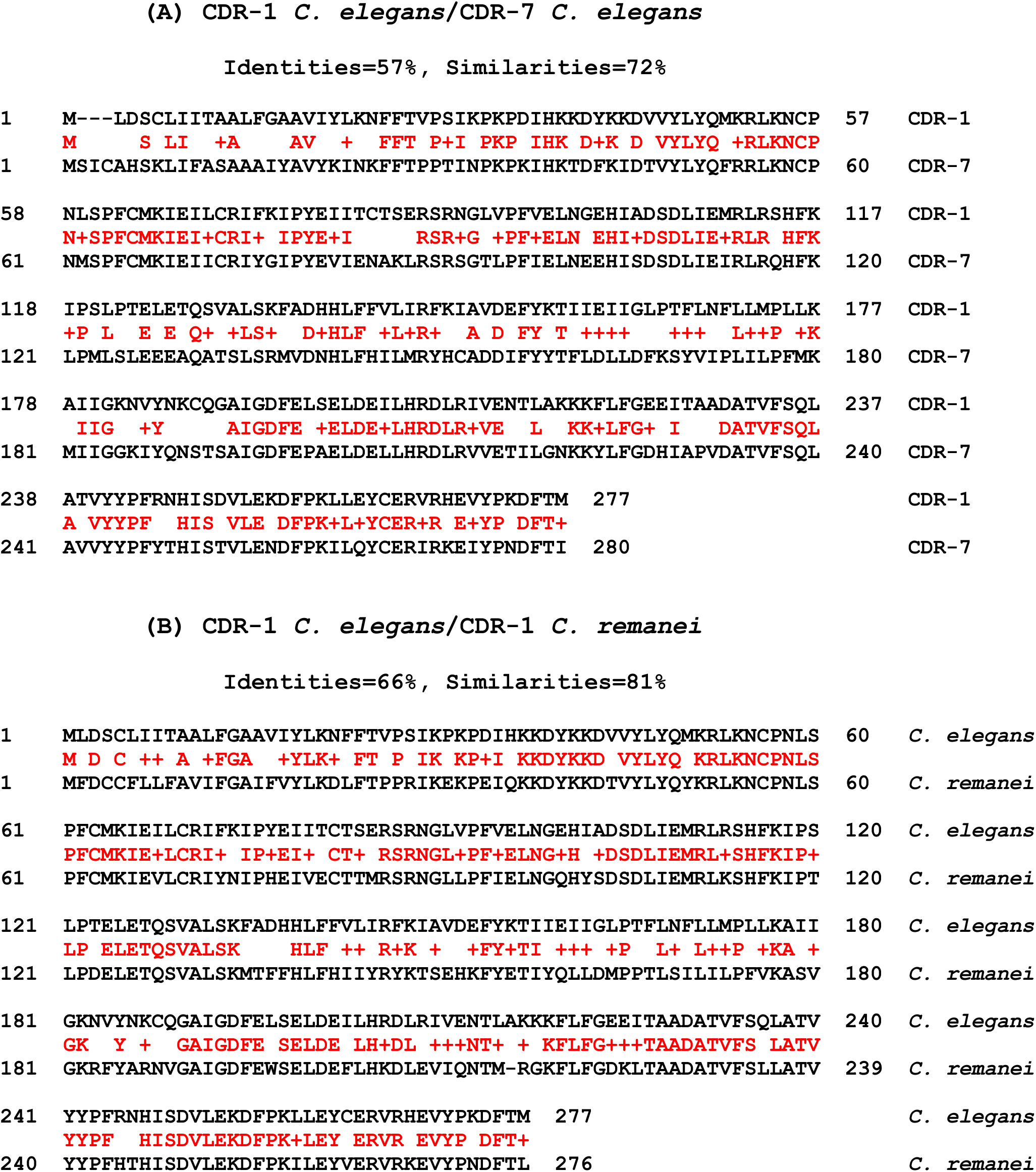
Amino acid sequence identities of the multiple CDR proteins of *C. elegans* and related species. (A) CDR-1 and CDR-7 proteins of *C. elegans* are aligned, and reveal high percentages of amino acid identities (57%) and similarities (72%). The alignment used NCBI Global Align. Comparable results were found in aligning different combinations of *C. elegans* CDR proteins. CDR proteins of *C. briggsae* and *C. remanei* also showed high percentages of amino acid identities and similarities comparing different proteins of the same *Caenorhabditis* species. (B) Alignment of the CDR-1 proteins of *C. elegans* and *C. remanei* exhibits a high level of homology, with 66% identical amino acids and 81% similar amino acids using NCBI Global Align. In addition, other alignments of CDR proteins of *C. elegans* and different *Caenorhabditis* species also displayed high levels of identical and similar amino acids.

For Figure 8, the same database was employed to determine which genes are adjacent to *cdr* and *faxc* genes.

**Figure 8.**
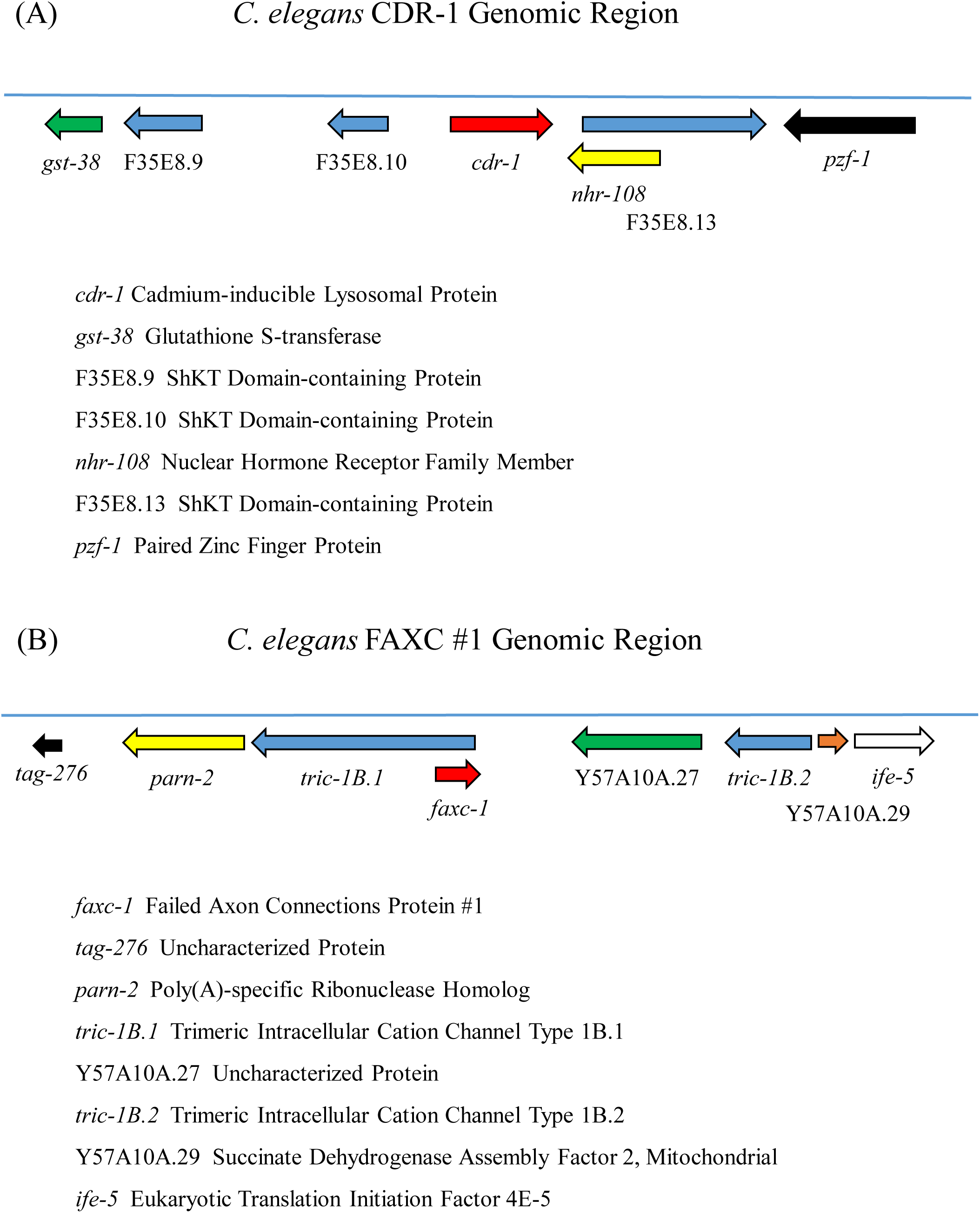
Adjacent genes of the *C. elegans cdr-1* gene and the *faxc-1* gene. The figure demonstrates that different neighboring genes flank the *cdr-1* and *faxc-1* genes. The immediate neighbors of the *cdr-1* gene (Figure 8A) are the genes for an ShKT domain-containing protein and a nuclear hormone receptor family member (*nhr-108*). In contrast, the genes next to the *faxc-1* gene (Figure 8B) are the genes for a trimeric intracellular cation channel protein and an uncharacterized protein. These differences in neighboring genes extend through the genomic regions shown in (A) and (B). The genomic regions of *cdr-1* and *faxc-1* are therefore very different. In addition, the *C. elegans* metaxin 1 gene *(mtx-1*) and metaxin 2 gene (*mtx-2*) have different neighboring genes compared to the *cdr* and *faxc* genes and to each other.

## 3. RESULTS AND DISCUSSION

### 3.0. Occurrence of CDR Genes

CDR (cadmium-responsive) genes were initially identified in the model invertebrate *C. elegans* (*Caenorhabditis elegans*), a nematode or roundworm (Liao et al., 2002). As shown in this report, genes homologous to CDR genes are also present in the related nematodes *C. briggsae* (genome sequence: Stein et al., 2003) and *C. remanei* (Fierst et al., 2015). In addition, CDR genes were detected in other nematodes such as *Necator americanus* (New World hookworm; Tang et al., 2014), *Haemonchus contortus* (barber’s pole worm), *Ancylostoma ceylanicum* (hookworm), *Angiostrongylus cantonensis* (rat lungworm), and *Diploscapter pachys*. In general, database searches indicated that CDR genes are only found in species of the phylum Nematoda. The genes were not detected in vertebrates, invertebrates other than nematodes, plants, protists, or fungi.

Unlike CDR proteins with their narrow distribution among species, metaxin and FAXC proteins, which have structural features in common with CDR proteins, are widely distributed among diverse vertebrates, invertebrates, and other organisms.

Perhaps surprisingly, some bacterial genomes have sequences that code for proteins with homology to CDR proteins (K.W. Adolph, unpublished). The bacterial proteins are about equally homologous to CDR and FAXC proteins, but there is little homology to metaxin proteins. Examples of these bacteria include the Gram-negative bacteria *Azotobacter salinestris* and *Pseudomonas hussainii*.

### 3.1. Protein Domain Structure of CDR Proteins

The presence of conserved protein domains, in particular the GST_N_Metaxin and GST_C_Metaxin domains, is an important structural feature in classifying proteins as metaxins. The metaxins of vertebrates and invertebrates, as well as the metaxin-like proteins found in plants, fungi, protists, and bacteria, all contain GST_N_Metaxin and GST_C_Metaxin domains. Moreover, these domains were found to be a characteristic feature of the FAXC proteins of vertebrates and invertebrates (Adolph, 2023).

As demonstrated in Figure 1, the CDR-1 protein of *C. elegans* also possesses GST_N_Metaxin and GST_C_Metaxin domains. The figure includes for comparison the GST_N_ and GST_C_Metaxin domains of *C. elegans* metaxin 1, metaxin 2, and FAXC #1. In addition, the Tom37 domain, also shown in the figure, is a major domain of metaxin and FAXC proteins. However, the Tom37 domain is not a characteristic domain of CDR proteins. Like many other invertebrates, *C. elegans* has multiple predicted FAXC proteins, with FAXC #1 included in the figure.

Figure 1 further shows that CDR proteins of several other nematode species possess the same conserved protein domains as *C. elegans*. These include *C. briggsae*, *C. remanei*, and *Necator americanus*. The GST_N_Metaxin and GST_C_Metaxin domains are the major conserved domains for each species, with no other major domains.

Multiple CDR genes are a prominent feature of the CDRs. For *C. elegans*, seven CDR genes have been detected (*cdr-1* through *cdr-7*), and both *C. briggsae* and *C. remanei* have multiple genes. Figure 1 includes the predicted CDR-1 protein of *C. elegans*, CDR-7 of *C. briggsae*, and CDR-5 of *C. remanei*. The conserved protein domain structures are similar for all of the multiple CDR proteins. [Genome regions that have multiple, adjacent CDR genes are shown in Figure 6 for *C. elegans* along with *C. briggsae* and *C. remanei*, and are discussed in Section 3.6.]

### 3.2. Conserved α-Helical Secondary Structure of CDR Proteins

Figure 2 reveals a striking conservation of α-helical segments for CDR proteins, metaxin proteins, and FAXC proteins. The CDR-1 proteins of *C. elegans* and other nematodes have secondary structures characterized by the same pattern of eight α-helices H1 – H8, plus an extra helix H0. This pattern of helices is also found for *C. elegans* CDR-2 through CDR-7 proteins, as well as the multiple CDR proteins of *C. briggsae* and *C. remanei*. The abundance of α-helical segments only leaves room for four short segments of β-strand, S1 through S4 (see Section 3.3). The pattern of α-helices labeled H1 – H8 in the figure was originally identified for the metaxins. In addition, metaxin 1 (HMTX1P in the figure) and also metaxin 3 have an extra helix H9 near the C-terminus, while metaxin 2 lacks H9 but has an extra helix near the N-terminus. FAXC proteins, in particular human FAXC isoform 1 in the figure, share the same pattern of α-helical segments H1 – H8 as the metaxins and CDR proteins. They also have extra helices near the N-terminus.

A feature of metaxin 1 proteins, and many metaxin-like proteins, is a transmembrane α-helical segment near the C-terminus. For example, human metaxin 1 has a transmembrane helix between residues 272 and 294 of the 317 amino acid protein. The helix could function to anchor the protein to the outer mitochondrial membrane, consistent with the role of metaxin 1 in protein import into mitochondria. CDR proteins, however, were not found to have a C-terminal transmembrane α-helical segment. This would indicate that being anchored like metaxin 1 to membranes is not integral to the functioning of CDR proteins. FAXC proteins also lack transmembrane α-helical segments.

### 3.3. Conserved Metaxin β-Sheet Motif: Presence in CDR Proteins

As demonstrated in Figure 2, the secondary structures of metaxin, FAXC, and CDR proteins are dominated by α-helical segments that are highly conserved. But as the figure shows, four, short β-strand segments are also found for human metaxin 1 (HMTX1P) and FAXC isoform 1, and for the CDR-1 proteins of *C*. *elegans* and other nematodes. The β-strand segments are labeled S1 through S4. For all the CDR-1 proteins, the β-strand segments are composed of 5 amino acids (S1 and S2) or 4 amino acids (S3 and S4). For the human metaxin 1 β-strand segments, S1 and S2 have 5 amino acids, S3 has 3, while S4 has 2. The β-strand segments for human FAXC isoform 1 in Figure 2 are all composed of 4 amino acids. Three of the β-strands, S2, S3, and S4, are located between α-helical segments H1 and H2, while β-strand S1 is N-terminal to H1.

In the 3D structures predicted by AlphaFold (Jumper et al., 2021), a characteristic β-sheet motif (Figure 3A,B) has been detected for CDR proteins, metaxin proteins, and FAXC proteins. Because metaxin genes are widely distributed among vertebrates and invertebrates compared to CDR genes, the motif is widely distributed and could be termed the metaxin β-sheet motif. In the 3D structures, the β-strand segments form a planar arrangement, with some distortion especially for S4 and also S3. The strands are in the order and general orientation shown in the figure: S2 (↑), S1 (↑), S3 (↓), and S4 (↑).

The metaxin β-sheet motif is present in the AlphaFold 3D structures of all the metaxin, FAXC, and CDR proteins that have been studied. In particular, *C*. *elegans* FAXCs and metaxins 1 and 2, in addition to *C*. *elegans* CDRs, possess the metaxin β-sheet motif. It is also found in the metaxin proteins of higher vertebrates including human metaxin 2 and metaxin 3. And the motif is in the FAXCs of fungi such as *Anaeromyces robustus* (Division: Neocallimastigomycota), *Spizellomyces punctatus* (Chytridiomycota), and *Heliocybe sulcata* (Basidiomycota).

Furthermore, the motif is present in the FAXC homologs of bacteria such as *Pseudomonas aeruginosa* and *Legionella jordanis*, both of the Gammaproteobacteria class.

### 3.4. Relationship of CDR, FAXC, and Metaxin Proteins: Phylogenetic Analysis

The phylogenetic results in Figure 4 demonstrate that *C. elegans* CDR proteins are closely related by evolution to FAXC proteins and to metaxin proteins. This is in keeping with the shared structural features of these proteins, in particular the conserved protein domains and patterns of α-helices and β-strands. CDR proteins 1 through 5 of *C. elegans* are shown in the figure, and form a separate group compared to the groups of FAXC and metaxin proteins. Four vertebrate proteins (human, mouse, *Xenopus laevis* frog, and *Danio rerio* zebrafish) make up the group of FAXC proteins and also the groups of metaxin 1, metaxin 2, and metaxin 3 proteins.

Five of the seven CDR proteins of *C. elegans* (CDR-1 through CDR-5) are included in the figure, and are seen to be more closely related to FAXC proteins than to metaxin proteins. Among the metaxin proteins, the metaxin 1 and metaxin 3 groups are more closely related compared to the metaxin 2 group. In agreement with these phylogenetic results, human metaxins 1 and 3 have a higher percentage of identical amino acids (45%) than metaxins 1 and 2 (22%). For other vertebrate species, the percentages of identical amino acids for metaxins 1 and 3 compared to metaxins 1 and 2 are similar to the percentages for the human metaxins.

### 3.5. Amino Acid Sequence Homology of CDR, Metaxin, and FAXC Proteins

The conserved protein domains (Figure 1), patterns of α-helical segments (Figure 2), and β-sheet structures (Figure 3) along with the phylogenetic results (Figure 4) show that CDR proteins, as well as FAXC proteins, have metaxin-like structural features. But alignments of amino acid sequences of CDR proteins with metaxin proteins and with FAXC proteins (Figure 5) indicate that the three proteins represent distinct protein categories. This is because the alignments show *low* percentages of identical and similar amino acids in comparing CDR proteins with metaxin and FAXC proteins. Consequently, CDR proteins and FAXC proteins are different categories of proteins, and not just different names for metaxin proteins. The structures (protein domains, α-helical and β-strand segments) of the metaxins are shared by CDR and FAXC proteins, and are highly conserved in contrast to the more divergent amino acid sequences of the three categories of proteins.

Figure 5A includes the alignment of CDR-1 and metaxin 1 of *C. elegans*. Only 17% amino acid identities are observed. In Figure 5B, CDR-1 and FAXC #1 of *C. elegans* are aligned, and reveal only 26% identities. Other alignments display similar low percentages. CDR-3 and metaxin 1 of *C. elegans* have 15% identical amino acids, while CDR-5 (also known as heme responsive protein hrg-2) and metaxin 1 have 18% identities. With *C. elegans* metaxin 2, CDR-1 shows 18% identities. Additional CDR and FAXC combinations also show low percentages of identical amino acids. CDR-3 and FAXC #1 have 29%, CDR-5 and FAXC #1 have 30%, and CDR-1 and FAXC #2 have 26%. The higher percentages of identical amino acids of CDR proteins with FAXC proteins compared to CDR proteins with metaxin proteins indicate that CDR proteins have greater homology to FAXC proteins.

### 3.6. Multiple CDR Genes of *Caenorhabditis* Species: Genomic Regions

As presented in Figure 6, CDR genes of *C. elegans* and related species can be found as multiple genes that occur in close proximity. For *C. elegans*, the figure (top) includes a closely spaced cluster of six CDR genes (cdr-2 through cdr-7) located on chromosome V. The *C. elegans* genome consists of five autosomes (I – V) and an X chromosome. Three CDR genes of *C. briggsae* (middle of figure) are adjacent on *C. briggsae* chromosome V, and two CDR genes of *C. remanei* (bottom) are also near each other.

In addition to multiple CDR genes, the presence of multiple FAXC genes is a common feature of invertebrates (Adolph, 2023). The invertebrate phyla with multiple FAXC genes include not only Nematoda (*C. elegans* has at least two FAXC genes), but Mollusca, Arthropoda, and other phyla. The sea anemone *Exaiptasia diaphana* (Cnidaria) has 13 or more, the lancelet *Branchiostoma floridae* (Chordata) has at least 10, and *Trichoplax adhaerens* (Placozoa) 12 or more.

A single FAXC gene is generally found in vertebrates. These single genes include the human FAXC gene (with three isoforms) and the mouse FAXC gene. However, the zebrafish *Danio rerio* and the frog *Xenopus laevis* each have two FAXC genes due to a genome duplication. The zebrafish has *faxca* on chromosome 4 and *faxcb* on chromosome 16. *Xenopus* has *faxc.L* on chromosome 5L and *faxc.S* on chromosome 5S.

Three metaxin genes (metaxins 1, 2, and 3) are typical of vertebrates, and two (metaxins 1 and 2) are generally found with invertebrates. Multiple metaxins are therefore the rule for vertebrates and invertebrates. In some cases, additional metaxins may be present. For example, the fruit fly *Drosophila melanogaster* has a single metaxin 1 homolog (CG9393) but two metaxin 2 homologs (CG5662 and CG8004), which have 43% identical amino acids.

### 3.7. Multiple CDR Proteins of *Caenorhabditis* Species: Amino Acid Sequence Homology

The predicted proteins of the multiple CDR genes of *C. elegans* and other nematodes show *high* percentages of identical amino acids in alignments of pairs of sequences. Figure 7A includes, for example, *C. elegans* CDR-1 and CDR-7 proteins. The amino acid identities are 57% and the similarities are 72%. In Figure 7B, CDR-1 of *C. elegans* and CDR-1 of *C. remanei* are aligned. Identical amino acids are 66% and similar amino acids, 81%. Other examples also indicate that these high values are typical percentages of identical amino acids when different CDR protein sequences are aligned. For *C. elegans*, CDR-1 and CDR-3 have 56% identical amino acids, CDR-3 and CDR-7 have 51%, and CDR-2 and CDR-4 have 62%.

The general conclusion is that the multiple CDR proteins of the same species or between different related species have a high degree of amino acid sequence homology. The high level of conservation of CDR protein sequences indicates that CDR proteins encoded by different genes, if expressed, would have similar structures. This suggests that the proteins are likely to function in a similar manner.

### 3.8. Neighboring Genes of CDR Genes

The genes adjacent to CDR genes of *C. elegans* differ from the genes adjacent to *C. elegans* FAXC and metaxin genes. Figure 8 displays in (A) the genes next to the *C. elegans cdr-1* gene, while (B) contains the genes next to the *C. elegans faxc-1* gene. The neighboring genes of the *cdr-1* and *faxc-1* genes are very different. The *cdr-1* gene on chromosome V is between the genes for an ShKT domain-containing protein and a nuclear hormone receptor family member (*nhr-108*). In contrast, the *C. elegans faxc-1* gene on chromosome II is between the genes for a trimeric intracellular cation channel protein, which it overlaps, and an uncharacterized protein. It should be noted that since *C. elegans* genes *cdr-2* to *cdr-7* belong to a *cdr* gene cluster on chromosome V (Figure 6), the neighboring genes for each of these *cdr* genes would include other *cdr* genes.

*C. elegans* metaxin genes are located in genomic regions that differ from both the *cdr* and *faxc* genomic regions. The metaxin 1 gene (*mtx-1*) on chromosome I is between the genes for transmembrane protein 107 (*tmem-107*) and an uncharacterized protein. The metaxin 2 gene (*mtx-2*) on chromosome III is flanked by genes for serine protease pcp-1 and ATP-dependent RNA helicase ZK686.2. There is therefore no similarity in the genomic regions of the *C. elegans cdr*, *faxc*, and *mtx* genes. For *C. briggsae* and *C. remanei*, and for other nematodes including *Necator americanus*. different genomic regions for *cdr*, *faxc*, and *mtx* genes were also found

The observation of different genomic contexts for CDR genes compared to metaxin and FAXC genes adds to the evidence in this report that CDR, metaxin, and FAXC proteins are related but distinct members of the metaxin and metaxin-like family of proteins.

## Notes

### Competing Interest Statement

The authors have declared no competing interest.

